# Bayesian multi-state multi-condition modeling of a protein structure based on X-ray crystallography data

**DOI:** 10.1101/2025.10.07.680994

**Authors:** Matthew Hancock, James Holton, James S. Fraser, Paul D. Adams, Andrej Sali

**Affiliations:** Department of Bioengineering and Therapeutic Sciences, University of California San Francisco, San Francisco, CA, USA; Department of Molecular Biophysics and Integrated Bioimaging, Lawrence Berkeley National Laboratory, Berkeley, CA, USA; Department of Biochemistry and Biophysics, University of California, San Francisco, CA, USA; Stanford Synchrotron Radiation Lightsource, SLAC National Accelerator Laboratory, Menlo Park, CA, USA; Department of Bioengineering, University of California Berkeley, Berkeley, CA, USA; Department of Pharmaceutical Chemistry, University of California San Francisco, San Francisco, CA, USA

## Abstract

An atomic structure model of a protein can be computed from a diffraction pattern of its crystal. While most crystallographic studies produce a single set of atomic coordinates, the billions of protein molecules in a crystal sample many conformational modes during data collection. As a result, a “multi-state” model that depicts these conformations could reproduce the X-ray data better than a single conformation, and thus likely be more accurate. Computing such a multistate model is challenging due to a lower data-to-parameter ratio than that for single-state modeling. To address this challenge, additional information could be considered, such as X-ray datasets collected for the same system under distinct experimental conditions (*eg*, temperature, ligands, mutations, and pressure). Here, we develop, benchmark, and illustrate MultiXray: Bayesian multi-state multi-condition modeling for X-ray crystallography. The input information is several X-ray datasets collected under distinct conditions and a molecular mechanics force field. The model consists of an independent coordinate set for each of several states and the weight of each state under each condition. A Bayesian posterior model density quantifies the match of the model with all X-ray datasets and the force field. A sample of models is drawn from the posterior model density using biased molecular dynamics (MD) simulations. We benchmark MultiXray on simulated CypA X-ray data. Using a second X-ray dataset improves the *R*_free_ from 0.105 to 0.089. We then demonstrate MultiXray on experimental temperature-dependent data for SARS-CoV-2 Mpro. Using multiple X-ray datasets improves *R*_free_ of the PDB-deposited structure from 0.253 to 0.237. MultiXray is implemented in our open-source *Integrative Modeling Platform* (IMP) software, relying on integration with *Phenix*, thus making it easily applicable to many studies.

## 2 Introduction

### A protein crystal is a heterogeneous mix of conformational states

X-ray crystallography has been a key experimental technique for determining structural models of proteins at atomic resolution [**Blundell Johnson 1976**]. A protein crystal typically contains between 10^6^ and 10^15^ protein molecules [RF96; Smi+15; WSF14]. These molecules exist in different conformations during data collection, because of dynamic and static disorder [KP90] [DBB04; Jen97; RP86][KPZ14; Kur+86]. Dynamic disorder results from approximately harmonic atomic motions within a local energy minima as well as anharmonic motions between the energy minima. Static disorder includes contributions from crystal imperfections and thermal inaccessibility of transition states between distinct conformations. In most crystallographic structure determinations, the conformational heterogeneity is modeled with isotropic B-factors, corresponding to a Gaussian distribution of conformations around the average conformation [Sun+19]. However, the B-factors generally cannot fully describe structural heterogeneity in a crystal [Hol+14; Vit+02]. A model that can better depict this heterogeneity will improve the satisfaction of X-ray data by the model. Such a model is also likely to be more informative; for example, it may better indicate an allosteric network and hidden cavities for small molecule binding (cryptic pockets) [BF15; HK07].

### Approaches to modeling a mixture of conformations

There are 2 approaches for computing models that depict multiple conformations from X-ray data. First, multiple conformations can be depicted by computing single conformation models, each one of which independently satisfies the X-ray dataset, as in the modeling of Nuclear Magnetic Resonance spectra [Sch97]. Such approaches may not find weakly occupied states in a reasonable amount of computation time, as they depend on overcoming potentially large barriers between local minima in the scoring function. Alternatively, a model can depict multiple conformations by introducing additional structural variables. For example, multiple conformations of a residue may be represented by distinct sets of atomic coordinates associated with alternative location identifiers and occupancies [WSF14]. All variables are then collectively fit against the X-ray data. The extent and detail of the additional degrees of freedom depend on the available computational power and the amount of data [Wan+24]. For example, qFit-3 avoids introducing excessive structural parameters by representing side chains and small backbone deviations as ensembles of one or more rotameric states [Ril+21]. Methods that refine multiple conformations have been limited to systems that diffract to better than 2.0Å resolution [RP86; WB00; Lev+07; Kur+91; BB94]. As a result, new methods are needed to accurately determine multiple conformations in a crystal based on X-ray diffraction data at worse than 2.0Å resolution.

### Multi-condition crystallography

Multiple X-ray datasets are often collected for the same system under distinct experimental conditions. One example is multi-temperature crystallography, where data collection is performed at temperatures from cryogenic (*eg*, 100 K) to near-physiological (*eg*, 310 K) [Tho23]. Protein dynamics within the protein crystal is highly dependent on temperature [FPT79; FSW91; TDP92]. Recently, models computed from higher temperature datasets have revealed a fuller set of conformations at atomic resolution [Fra+09; Hal04; Kee+14]. Comparisons of models of the same system computed at distinct temperatures show similar conformations, but with different relative stability [Ebr+21; Fra+09; Du+23]. Therefore, multiple X-ray datasets under distinct conditions contain mutual structural information and may increase the data-to-parameter ratio for informing a model depicting multiple conformations simultaneously.

### Computing a multi-state multi-condition model

Here, we seek to compute a model depicting multiple conformations (multi-state model) from multiple X-ray diffraction patterns collected under distinct conditions (multi-condition model) (Figure 1). As a result, we may be able to compute a multi-state model using multiple X-ray datasets without the need for ultra-high-resolution X-ray data. Modeling accommodates the varying stability of individual conformations at distinct experimental conditions by computing each state”s weight under each condition, while assuming the set of conformations is preserved across all experimental conditions. This model representation allows a single multi-state model to be informed by all X-ray datasets, thus significantly improving the data-to-parameter ratio for modeling. The assumption of a fixed set of conformational states under all conditions is not limiting, because conformations absent under some conditions can simply have a zero weight. To convert the input X-ray datasets into a model, we formulate a Bayesian posterior model density for the multi-state multi-condition model. A sample from the Bayesian posterior model density is drawn using biased molecular dynamics simulations where all states are jointly restrained by the satisfaction of all X-ray datasets. The method is benchmarked using synthetic X-ray datasets simulated from multi-state CypA structural models. As expected, the results demonstrate that multi-state multi-condition models are better in satisfying the data than either the single-state or single-condition models, given the structural sampling method used. We also illustrate the method by application to a real-world problem, corresponding to computing a multi-state model of the SARS-CoV-2 viral target M^pro^ from experimentally-determined multi-temperature X-ray datasets.

**Figure 1.**
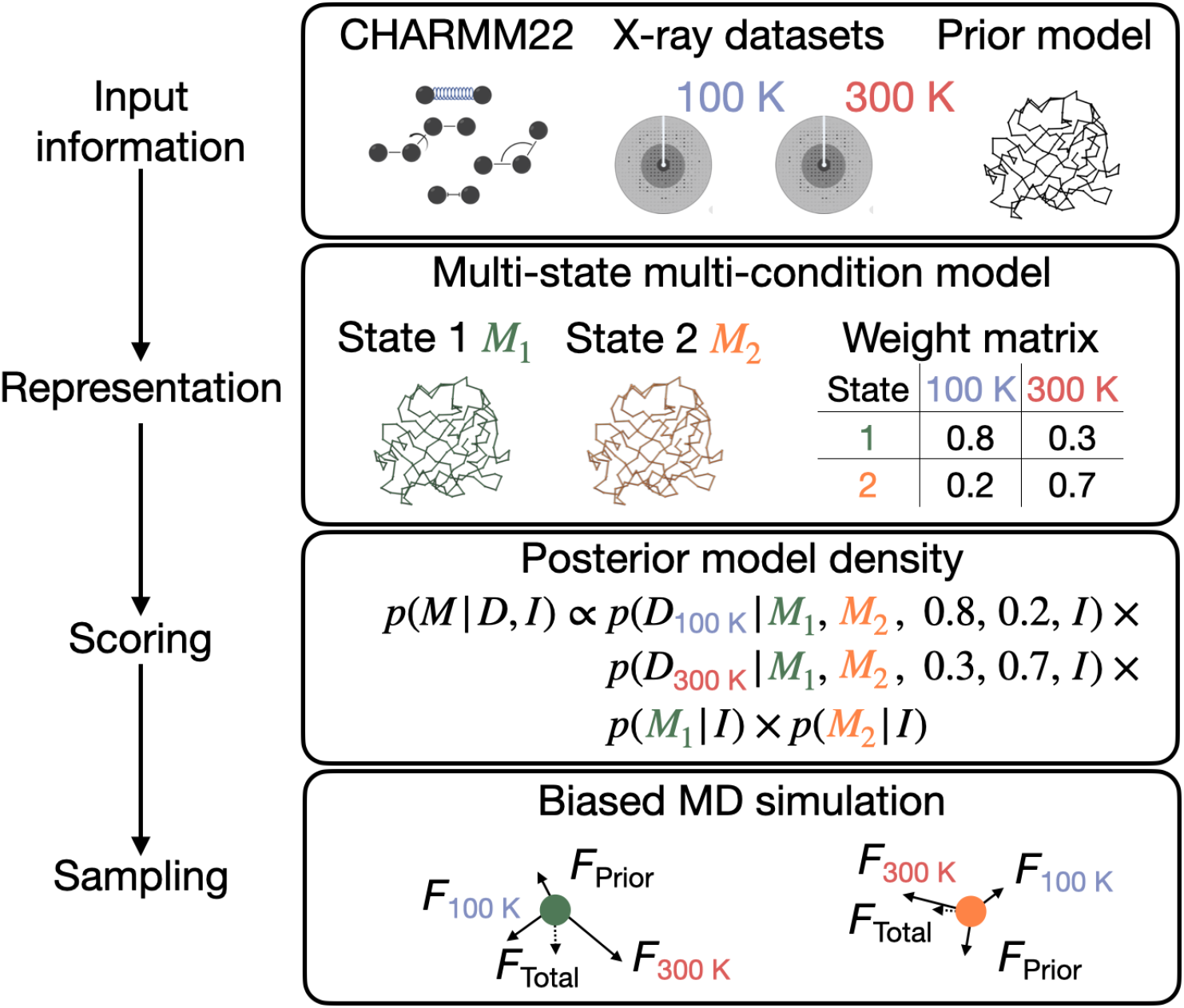
Four stages of modeling. Modeling is framed as a search for models whose computed properties match input information. Here, the input information is the CHARMM22 force field parameters, multiple X-ray datasets collected for the same system under distinct experimental conditions (*eg*, temperature), and a prior model. The model representation includes variables specifying atomic coordinates for each conformational state, along with a matrix of weight variables for each state under each condition. All states and weights are scored collectively for each X-ray dataset. Each state is individually scored against the molecular mechanics force field. A sample is drawn from the posterior model density using biased molecular dynamics simulations. All states are initialized by a prior model and the force on the atoms is computed from the satisfaction of all X-ray datasets and potential energy.

## 3 Methods

### Modeling approach

A model is a depiction of our knowledge about a system or process. We wish to inform the model based on the input information, which generally includes experimental data and prior models (*ie*, physical theories, statistical preferences, and other prior models). A model can then rationalize past observations and predict future ones. Modeling is the search for a set of models consistent with the input information. We aim to find all models that satisfy the input information, reflecting the uncertainty of the input information and the modeling process. In statistical modeling [**ref**], it is convenient to describe modeling in terms of its three aspects: (i) specifying all model variables (representation), (ii) ranking alternative models by their agreement with the input information (scoring), and (iii) generating a sample of good-scoring models (sampling). In addition, a model should be validated before being interpreted; for example, by examining the satisfaction of input information withheld from the modeling process. Multiple iterations of gathering input information, modeling, and validation are often necessary to compute a sufficiently precise model that satisfies input information [RS19; Sal21]. The following sections provide a detailed description of input information, representation, scoring, and sampling for our multi-state multi-condition modeling method.

### Input information

The input information may be used to inform any step of modeling: representation, scoring, and sampling. Here, we use the following three types of input information: First, X-ray datasets *D*_1_, …, *D*_*J*_ collected under distinct experimental conditions (*eg*, at distinct temperatures). Each X-ray dataset is a set of observed structure factor amplitudes 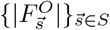 indexed by a scattering vector 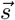. The X-ray datasets are used in scoring. Second, a starting atomic structure model; for example, a standard atomic structure model based on one of the X-ray datasets. This starting model informs both representation and sampling. Third, the CHARMM22 force field parameters [Bro+09]. The force field parameters are used in scoring.

### Representation

The representation defines the variables whose values are to be determined by modeling. The multi-state multi-condition model *M* includes the following three sets of variables: First, *N* states *M*_*i*_, where *N* is selected before modeling. Each state *M*_*i*_ is an independently parameterized model of a protein structure. The number and type of atoms (composition) of each state are based on the input model (Input information). Model representation in general depends on the input information and the question to be answered. Here, we include all protein atoms (including hydrogen atoms). The representation may also be expanded to include additional atoms/molecules (*eg*, solvent, ions, and ligands). Because we are computing a model depicting a set of discrete conformational states, all atomic B-factors are set to 15 and atomic occupancies to 1, although they could in principle also be sampled. Second, weight matrix *W*_*N×J*_ containing the weights of *M*_*i*_ under conditions *j, w*_*ij*_; the sum of the state weights under each condition (*ie*, columns) is 1. Third, nuisance variables, taken from established modeling methods, to improve the fit of the X-ray datasets by the model [Afo+13; LAU02]; these nuisance variables include 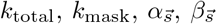, and variables describing the bulk solvent model. For clarity, we do not include nuisance parameters in our notation of a model.

### Scoring

A scoring function assesses the match of a model to the input information. Bayesian inference is probably the most rigorous approach to assess this fit. In Bayesian inference, the posterior model density *p*(*M* | *D, I*) describes the relationship between model *M*, prior information *I*, and data *D*. According to Bayesian inference, the posterior model density is factored into a prior *p*(*M*|*I*) and a likelihood *p*(*D*|*M, I*) [Mce15]:

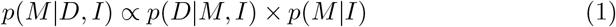

where *p*(*M* | *I*) is the probability density of a model given prior information and *p*(*D* | *M, I*) is the probability density of data given model and input information. It is often helpful to decompose the likelihood into a forward model *f* (*M*) that simulates a noiseless data observation and a noise model *N* (*f* (*M*); *D, σ*) that quantifies the difference between the observed data and data computed from the model [RS19].

#### Likelihood

Here, *D* is all *J* X-ray datasets. The likelihood of observing all X-ray datasets is factored into the independent likelihoods of observing each X-ray dataset:

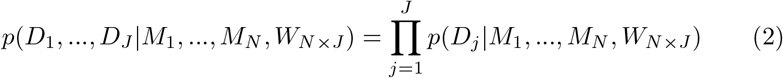

The likelihood for observing an X-ray dataset given a model depends only on the set of states and the weights of the states under the corresponding condition (matrix column):

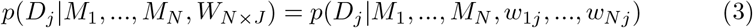

The likelihood for observing an X-ray dataset is further factored into the independent likelihoods of observing each structure factor amplitude 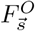 [LAU02]:

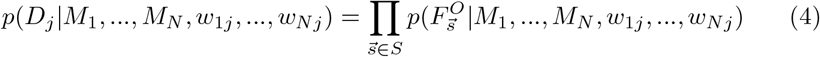

The likelihood of observing a structure factor amplitude is a noise model that quantifies the difference between the observed structure factor amplitude 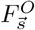 and the model structure factor amplitude 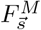 computed from the set of states and their weights [LAU02]:

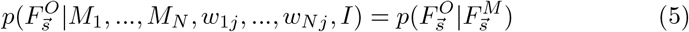

The noise model assumes that the real and imaginary parts of a complex structure factor **F** are sampled from a two-dimensional Gaussian distribution with variance 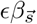 and scale parameter 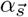 [LAU02]:

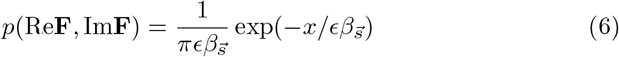

where:

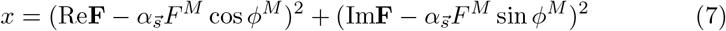

The likelihood for a structure factor amplitude is [LAU02]:

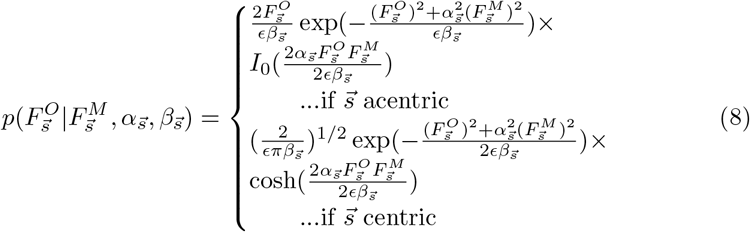

where *I*_0_ is the hyperbolic Bessel function of the first kind (*α* = 0) and cosh is the hyperbolic cosine function.

#### Forward model

The forward model 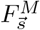 is the magnitude of the complex model structure factor 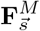:

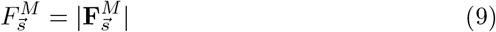

The model structure factor is the sum of the protein 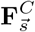 and bulk solvent structure factor 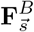 with an overall scale factor *k*_total_ and bulk solvent scaling factor *k*_mask_ [Lie+19].

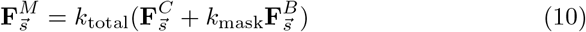

Computing the scattering of a crystal from a model of a protein structure assuming a perfect crystal is well established [Gui63]. Here, we aim to compute the scattering of a model of a protein structure with multiple states and their weights, representing conformational variability over space and time. Importantly, due to the nature of the crystallographic forward model, different structures and positions of molecules in a crystal can result in the same X-ray diffraction data, even when the data are noiseless, complete, and collected instantaneously. More specifically, the X-ray data is degenerate in the sense that it does not discriminate among the 5 types of crystal configurations enumerated in Figure 2. The degeneracy of a heterogeneous crystal and the weighted sum of the corresponding homogeneous crystals (Figure 2) enables the scattering of a crystal composed of *N* states to be computed from the weighted sum of the scattering of a perfect crystal containing each state. Thus, the structure factor for a multi-state model may be computed from the single state structure factor 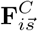 and the weight of the state in the heterogeneous crystal *w*_*i*_:

**Figure 2.**
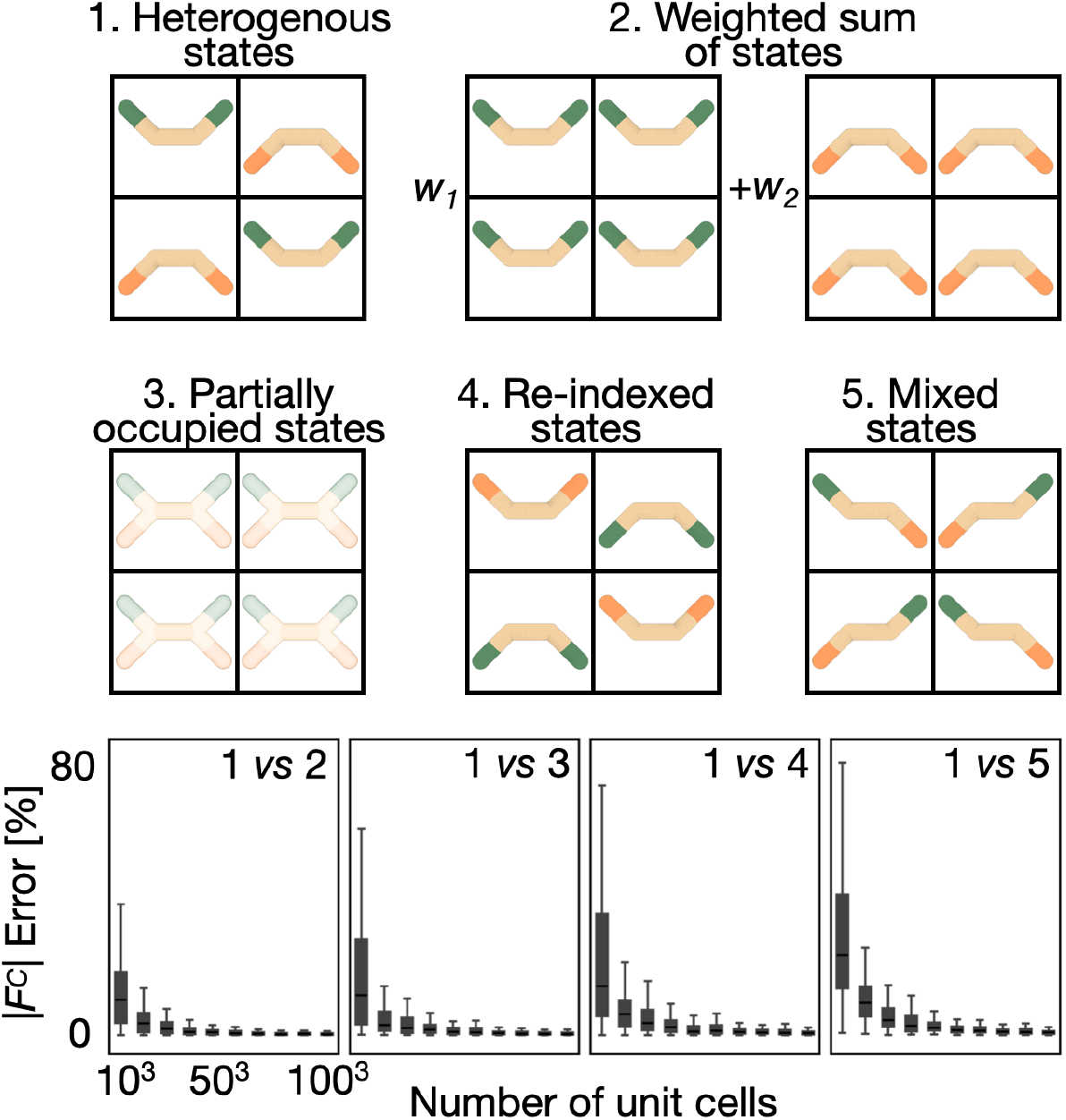
Degenerate scattering based on a multi-state model. The X-ray forward model will produce identical computed data, and therefore posterior model density, for the following 5 combinations of 2-state models and unit cells: (1) A single crystal with different unit cells containing atoms in either *M*_1_ or *M*_2_ with probability *w*_1_ and *w*_2_, respectively. (2) Two crystals, with the first crystal containing atoms in *M*_1_ and the second crystal containing atoms in *M*_2_, with the proportion of atoms in *M*_1_ and *M*_2_ corresponding to *w*_1_ and *w*_2_, respectively. (3) A single crystal containing identical partially occupied unit cells with atoms in *M*_1_ and *M*_2_ with occupancy *w*_1_ and *w*_2_, respectively. (4) The same as (1) but with the two state indices swapped. (5) A single crystal with different unit cells, each one of which contains a mixture of atoms in the two states in equal proportion. As the number of unit cells approaches infinity, the reciprocal lattice points simulated from a crystal in (2), (3), (4), and (5) converge to those for the crystal in (1). We simulate scattering for (1)-(5) using discrete Fourier Transform (DFT). As the number of unit cells in the crystal increases, the error between the reciprocal lattice points of (2)-(5) and (1) converges to The error is defined as the Euclidean distance between the structure factor and the reference structure factor, normalized by the magnitude of the reference structure factor.

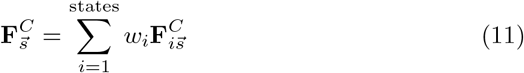

The degeneracy of a heterogeneous crystal and intermixed heterogeneous crystal (panels 1 and 5 in Figure 2) complicates the interpretation of a match between a multi-state model and an X-ray dataset. For example, without exhaustive sampling of all possible *N* -state models with *N* rotamers for residues A and B each, we do not know if the rotamer pair always occurs in the same molecule or is one of many possible combinations of the rotameric states in multiple molecules; in other words, we cannot rule out any combination of the rotameric states of A and B in one molecule.

As an aside, the degeneracy of the X-ray data might be broken by using additional information for modeling; for example, a potential energy function may rule out combinations of rotameric states that result in overlapping atoms. Additional sources of X-ray scattering, such as diffuse scattering, may also be used to break the degeneracy of Bragg data [WWF18].

Finally, 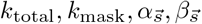 and 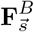 are optimized to maximize the similarity of all 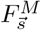 and 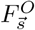 as described previously [Afo+13].

#### Prior

The prior for the multi-state multi-condition model is the product of a prior for each state:

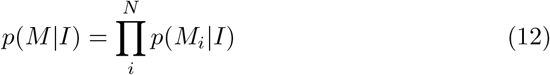

The prior for a state is the Boltzmann distribution corresponding to the potential energy of the state, which in turn includes the terms for bond lengths *b*, bond angles *θ*, dihedral angles *ϕ*, improper dihedral angles *w*, and non-bonded interactions *r*_*ij*_:

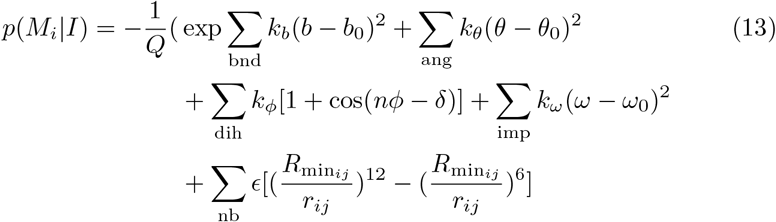

where the parameters *k*_*b*_, *b*_0_, *k*_*θ*_, *θ*_0_, *k*_*ϕ*_, *δ, k*_*w*_, *w*_0_, *ϵ*, and 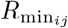 are obtained from the CHARMM22 molecular mechanics force field [Bro+09] and *Q* is the partition function, which is ignored in practice because it does not change the ranking of alternative models based on the potential energy function alone. Electrostatic interactions are ignored due to technical challenges.

#### Scoring function

To improve the numerical robustness, a Bayesian scoring function for ranking alternative models is the negative logarithm of the posterior model density:

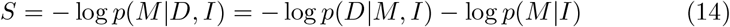

### Sampling

The purpose of sampling is to find all models consistent with the input information. In Bayesian modeling, this goal is achieved by computing a sufficiently converged estimate of *p*(*M* | *D, I*). To obtain the corresponding sample, we use biased molecular dynamics simulations to sample structural variables and random sampling to sample the weights of all states under all conditions. The molecular dynamics simulations are biased because the force on each atom reflects the entire scoring function, not just the potential energy. Thus, the force on each atom in each state, 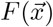, depends on all X-ray datasets as well as the potential energy of that state:

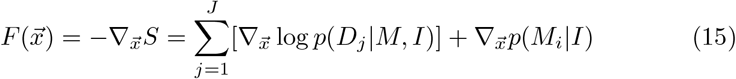

To calculate derivative of the likelihood more efficiently, a quadratic approximation of 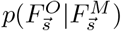 is used [LAU02].

We noticed that the magnitude of the prior gradient is much larger than that of the likelihood gradient. Thus, to facilitate more efficient sampling, we introduced 2 scaling parameters *w*_xray_ and *w*_auto_ in the calculation of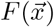:

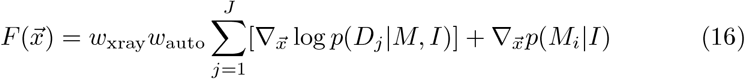

*w*_auto_ is computed automatically during sampling, by calculating the average magnitude of the likelihood gradient for all atoms and scaling it to be equal to the average magnitude of the prior gradient for all atoms [BB94]. *w*_xray_ reflects an empirical observation that slightly under-weighting the likelihood gradients relative to the prior gradients improves the exploration of the search space (as measured by the *R*_free_ of the best-scoring model in the sample). We generally compute our sample with multiple values of *w*_xray_ (*eg, w*_xray_ = 1.0, 0.5, 0.25).

The atomic positions of all states are initialized to the input model (Input information). The “velocity” of all atomic coordinates is sampled from a “Boltzmann distribution” with “temperature” *T* = 5000*K*. A high sampling “temperature” allows exploration of a rugged multi-state posterior model density. Because of the likelihood, the force on the atoms is non-conservative, and a thermostat is used to maintain a simulation temperature of *T* [Bur+12]. We presently do not include explicit solvent or ligand molecules in the simulations, although there is no principled reason preventing the inclusion of such molecules in the future; instead, we rely on the likelihood to provide accurate models. As the last step of our sampling workflow, the atomic positions of models in the sample are refined with a conjugate gradients method (*w*_xray_ = 1), resulting in models at local minima of the scoring function.

### Software availability

The software, input files, and output files are freely available at as part of our open-source *Integrative Modeling Platform* (IMP) (https://integrativemodeling.org/2.20.0/doc/manual/) [Rus+12]. IMP relies on an interface with *Phenix* (https://phenix-online.org/) [Lie+19], as described previously [Han+22].

## 4 Results

### Framework for analyzing a modeling method

If the ground truth (native) is known, a modeling method may be evaluated by its ability to find the native as the best-scoring model. If models could be enumerated with sufficient precision, the only feature of the scoring function required for accurate modeling would be that its global minimum corresponds to the native (global minimum accuracy). However, in practice, models cannot be enumerated with sufficient precision due to the high dimensionality of the search space. Therefore, we use stochastic search methods to find the global minimum. The efficacy of most search methods depends strongly on the scoring function having a funnel shape around the global minimum. The radius of convergence quantifies how far from the global minimum does the funnel assist the sampling convergence to the global minimum (the width of the funnel). The smoothness of the funnel is defined as the correlation between the score and model coordinates. Most sampling methods benefit from a smoother funnel with a larger radius of convergence.

We benchmark our modeling method by analyzing the 2-state 2-condition scoring function. Because the prior has been evaluated previously, we focus on the negative log-likelihood and refer to the negative log-likelihood as the scoring function. First, we estimate the global minimum of the scoring function. Second, we quantify the thoroughness of the search process by plotting the best score found as a function of the amount of sampling (convergence curve). Third, we interpret the differences between these plots for different scoring functions by considering their radius of convergences and smoothness. If the differences in the convergence curve cannot be explained by the differences in the radius of convergence and smoothness of the scoring function, then they must be a result of higher-dimensional characteristics of the landscape.

### Native model

There is no large set of multi-state multi-condition models with corresponding experimentally measured X-ray datasets. Thus, we use a synthetic benchmark, where a reflection dataset is computed, rather than measured, from a native. The benchmark contains a 2-state 2-condition model constructed, in part, from a previously determined CypA structure [Fra+09; Kee+15].

### Native reflection datasets

For a native, X-ray datasets *D*_1_ and *D*_2_, corresponding to condition 1 and condition 2, are simulated via the forward model at 2.0Å resolution based on the experimentally determined experimental unit cell dimensions and space group [Ebr+21] (Figure 3). Gaussian noise is applied to the simulated structure factor amplitudes. Reflections within each resolution shell are randomly withheld for model assessment. The datasets are highly correlated and thus the same set of reflections is withheld from both dataset 1 and dataset 2.

**Figure 3.**
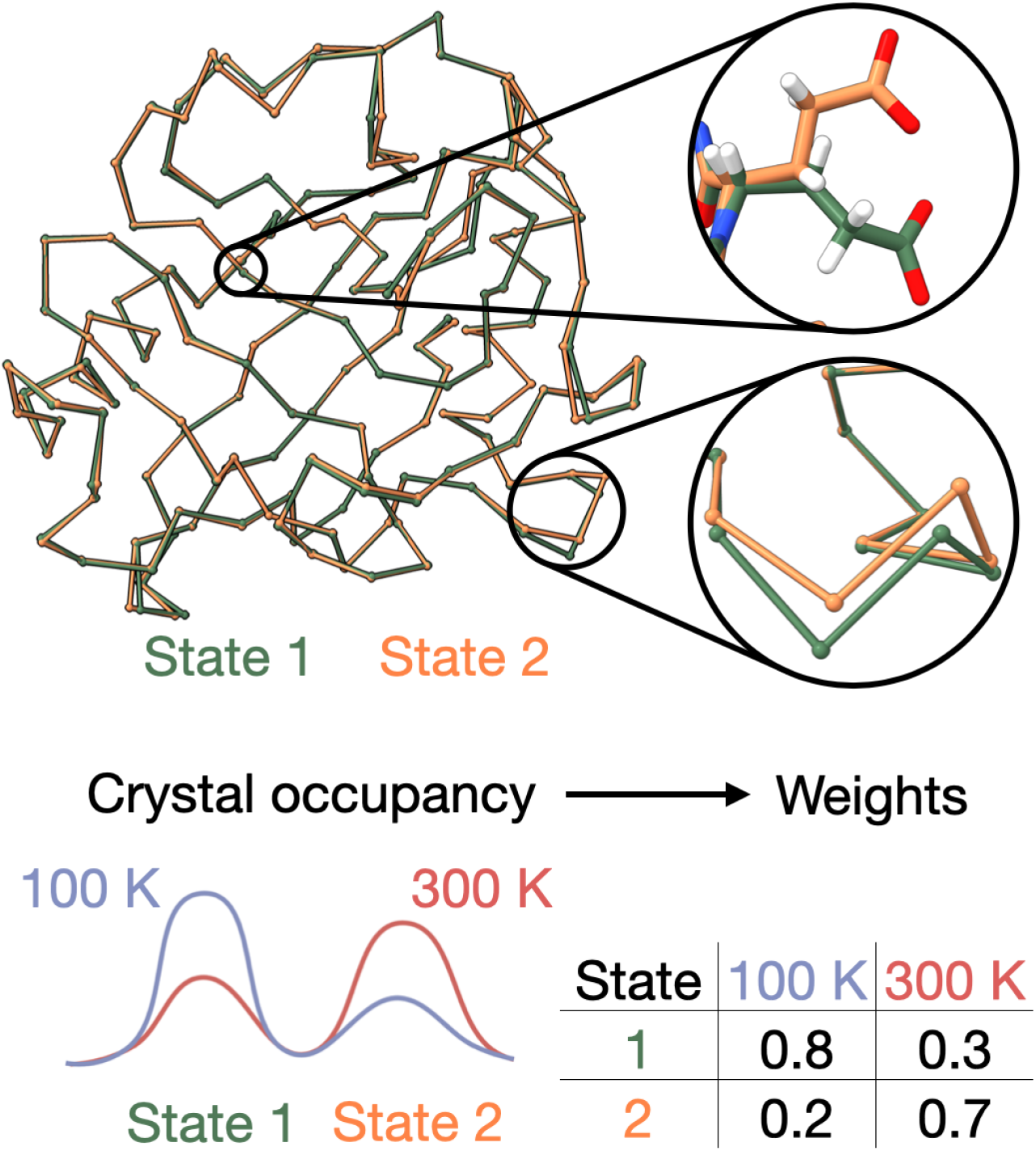
Native model for benchmarking. A 2-state 2-condition native model based in part on a previously determined structure model of CypA (PDB: 3K0M). The native contains 2 conformational states shown in green and orange, respectively. The 2 states exhibit structural heterogeneity typical of an actual crystal: small backbone deviations and large side chain deviations. The population of each state under the 2 simulated conditions varies. State 1, representative of a low energy state, is the dominant state at 100 K while state 2, representative of a high energy state, is the dominant state at 300 K. The corresponding weight matrix is shown.

### Model difference and accuracy

We define the model difference of a *N*_*A*_-state *J* -condition model by its deviation from a *N*_*B*_-state *J* -condition model where all states are compositionally identical as follows:

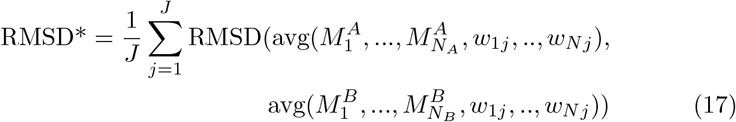

The inside sum is the RMSD between the weighted average of *M*^*A*^ and the weighted average of *M*^*B*^ using the *j* column of the respective weight matrices. In the weighted average, the atomic position vector *k* is the weighted sum of the respective atomic position vectors across all states.

Computing the model error in this fashion is useful for the following reasons. First, we can compute the model error when *N*_*A*_ ≠ *N*_*B*_. Second, if *N*_*A*_ = *N*_*B*_ = 1, then the model error is the average of the RMSD function across all conditions.

Third, the model error equals 0, if and only if the atomic position vector of the model average of *M*^*A*^ and the model average of *M*^*B*^ are equal for all atoms for all conditions. It is possible, however, for distinct models to have the same weighted average structure and thus an error of 0. Fourth, the model error shares the symmetry properties with the multi-state X-ray forward model for a given condition. The model error and multi-state X-ray forward model are a function of the set of atomic positions for a given atom *k* (along with their weights) and, therefore, independent of the assignment of an atom to a specific state. For example, the model error would not change, if the states in *M*^*A*^ or *M*^*B*^ are re-indexed. More generally, the model error would not change, if the positions of atoms *k* in any 2 states are swapped (state mixing).

### Accuracy of the global minimum

Because the benchmark is performed with data simulated from the native, we assume with high confidence that the native is the global minimum of the scoring function. Though it is difficult to prove this assumption rigorously (*eg*, by enumeration), there is no decoy model out of a set of 1000 decoy models that better satisfies the data than the native (Figure 4).

**Figure 4.**
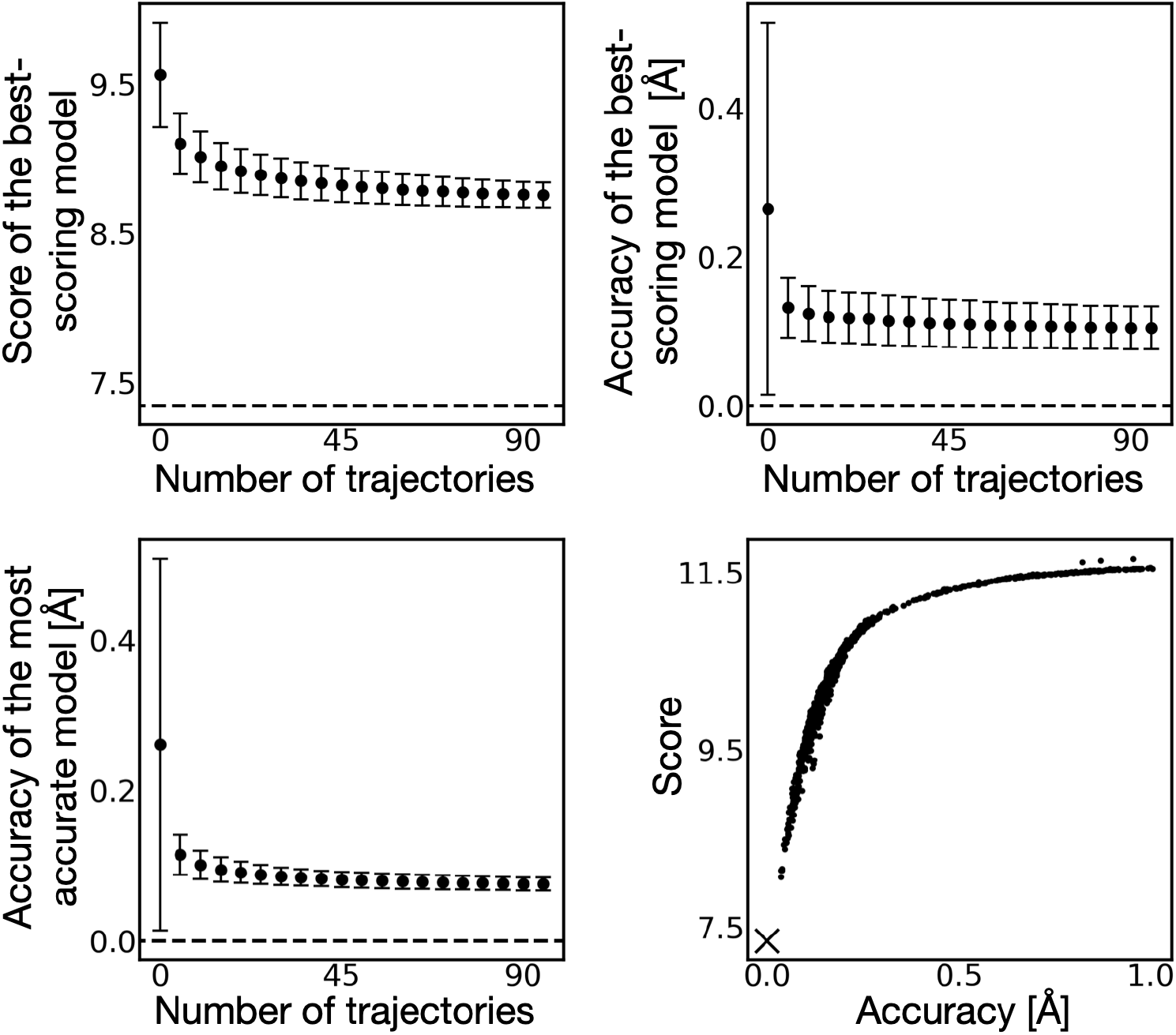
Benchmark of multi-state multi-condition modeling. Top left panel shows the mean and standard deviation of the score (joint log-likelihood) of the best-scoring model from 1000 sub-samples consisting of an increasing number of MD trajectories. The score of the native is shown as a dashed line. Top right panel shows the mean and standard deviation of the accuracy of the best-scoring model for the same sub-samples. Bottom left panel shows the mean and standard deviation of the accuracy of the most accurate model for the same sub-samples. Bottom right panel shows the score (joint log-likelihood) and accuracy of 1000 decoy 2-state 2-condition models generated in 5000 short MD simulations, selected randomly to approximately evenly span an accuracy range from 0 to 1Å. X indicates the score and accuracy of the native model.

### Sampling convergence

To quantify the thoroughness of the search process, we plot the sampling convergence of the scoring function (Figure 4). The score of the best-scoring model, the accuracy of the best-scoring model, and the accuracy of the most accurate model found in the sample all converge, suggesting modeling would not benefit significantly from additional sampling. However, as expected for a stochastic search, it does not find the global minimum of the scoring function, instead becoming stuck in local minima near the global minimum.

To show the advantage of considering an addition X-ray dataset, we compare the best-scoring 2-condition model to the best-scoring 1-condition model, as follows. We report the best *D*_1_ *R*_free_ and the best *D*_2_ *R*_free_ from the sample of *P* (*M*_22_|*D*_1_, *D*_2_, *I*). Next, we compute two 1-condition samples from *p*(*M*_21_|*D*_1_, *I*) and *P* (*M*_21_|*D*_2_, *I*). We report the best *D*_1_ *R*_free_ from the sample of *p*(*M*_21_|*D*_1_, *I*) and the best *D*_2_ *R*_free_ from the sample of *p*(*M*_21_|*D*_2_, *I*) (Table 1). Finally, we report the corresponding *R*_work_ for all models (Table 2). A 2-condition model improves the *R*_free_ from 0.019 to 0.0197 and the *R*_free_ for condition 2 from 0.105 to 0.089. The improved recovery of the native by modeling using data from 2 conditions suggests there are improvements to scoring and/or sampling relative to modeling using data from a single condition. For condition 1, we show that the best-scoring 2-condition model has significant improvements to the recovery of the native backbone and side chains relative to a 1-condition model (Figure 5).

**Table 1:** Benchmark results. The *R*_free_ of the best-scoring model for each dataset when 1 and 2 X-ray datasets are used in modeling. The best-scoring model for a dataset is the sample model that best satisfies the free reflections of a dataset. The same set of free reflections has been used for all datasets.

| Model representation | | $R_{\text{free}}$ | |
| --- | --- | --- | --- |
| States | Cond. | 1 (2.00 Å) | 2 (2.00 Å) |
| 2 | 1 | 0.115 | 0.135 |
| 2 | 2 | 0.136 | 0.102 |

**Table 2:** Benchmark results. The corresponding *R*_work_ for all entries in previous table.

| Model representation | | $R_{\text{work}}$ | |
| --- | --- | --- | --- |
| States | Cond. | 1 (2.00 Å) | 2 (2.00 Å) |
| 2 | 1 | 0.109 | 0.105 |
| 2 | 2 | 0.097 | 0.089 |

**Figure 5.**
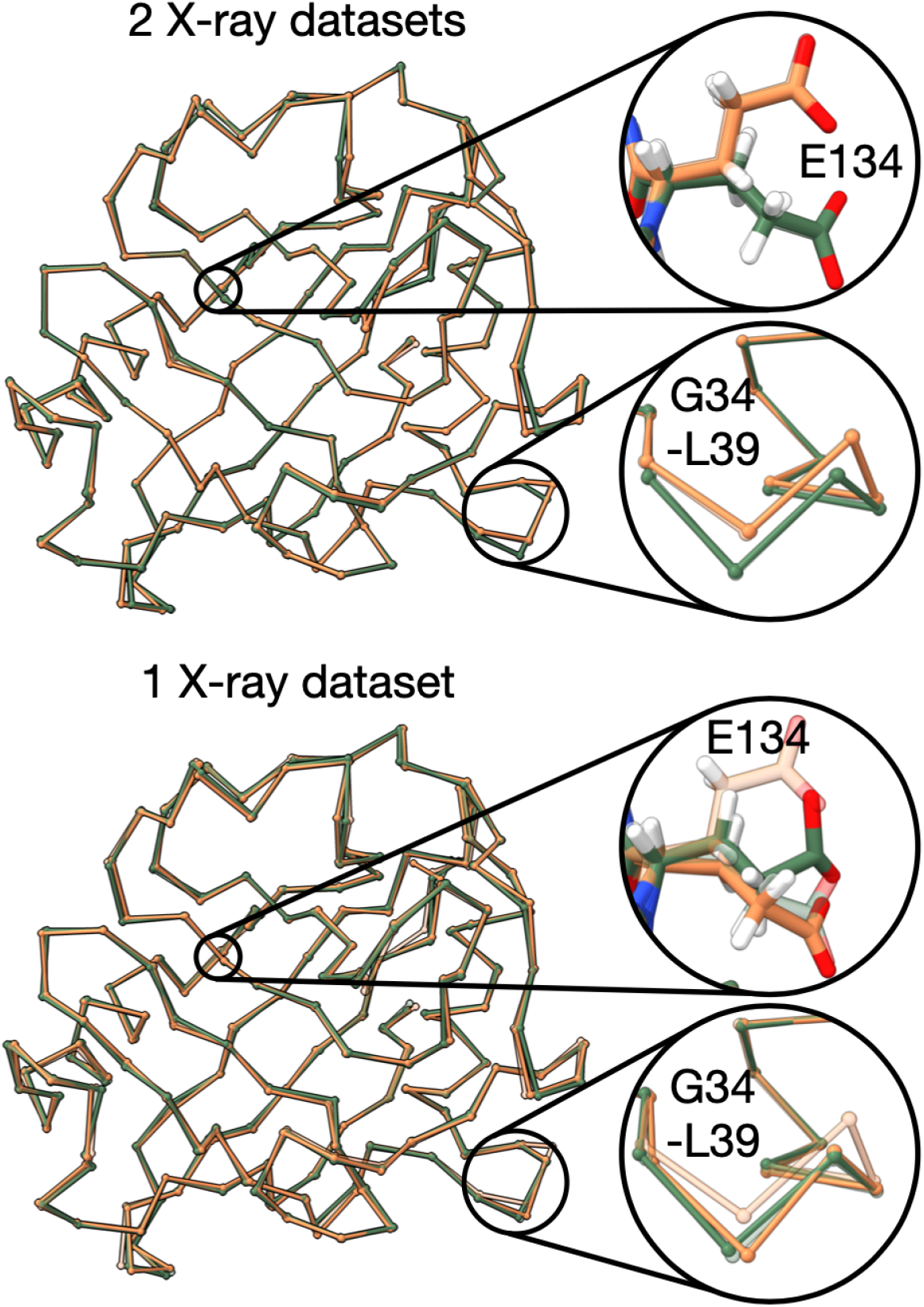
Using data from multiple conditions improves recovery of native. Top, the best-scoring 2-state 2-condition CypA model for condition 1, as identified by the best *R*_free_ for dataset 1. The model is overlayed over state 1 and state 2 of the native (transparent). To illustrate the accuracy of the backbone and side chain model heterogeneity, the Glu134 atoms and the C*α* atoms of Leu39 - Gly47 are shown. The 2-condition model correctly fits both the backbone and side chain heterogeneity of the native. Bottom, the best-scoring 1-state 1-condition model for condition 1, as identified by the best *R*_free_ for dataset 1. The 1-condition model does not correctly fit the side chain and backbone heterogeneity of the native. Both states 1 and 2 of the 1-condition model are a closer fit to state 1 of the native than state 2 of the native.

### The shape of the scoring function

The convergence of the model search to the global minimum of the 1-state scoring function (Figure 4) suggests that the scoring function is smooth and has a large enough radius of convergence to facilitate the model search by molecular dynamics simulations, given the initial conditions.

The joint likelihood and the accuracy are a function of both the weights and conformations of the states. We show the scoring function with 2 additional decoy sets (Figure 4). The first decoy sets contain 2-state 2-condition models where the conformations are the same as the native, but the weights are randomized. The second decoy set contains 2-state 2-condition decoy models where the conformations are randomized, but the weights are the same as that of the native. When only conformations are varied, the score is strongly correlated to the accuracy. When only the weights are varied, the score is more weakly correlated to the accuracy.

### Multi-state multi-condition models of SARS-CoV-2 Mpro

We illustrate our method by computing multi-state multi-condition models of the SARS-CoV-2 main protease (M^pro^), a therapeutic target [Jin+20]. Data was collected and modeled previously at 100 K, 240 K, 277 K, 298 K, 298 K* (high humidity), and 310 K [Ebr+21]. These datasets, as deposited in the PDB, include a distinct set of free reflections. Because all X-ray datasets are highly correlated, we create a new set of free reflections used for validating the model against all X-ray datasets, maintaining the proportion of free and work reflections in each resolution bin. Using the same set of free reflections across all dataset, ensures no information contamination between the free and work reflections of highly correlated X-ray datasets.

For each subset of 3-datasets (20 in total), 2 datasets (15 in total), and 1 dataset (6 in total) as input, we compute a sample of 2-state (3/2/1)-condition models.

For each X-ray dataset, we report the *R*_free_ of the best-scoring 1-state 1-condition, 2-state (3/2/1)-condition model (Figure 6, Table 3), and the corresponding *R*_work_ (Table 4). In all 6 cases, an additional state (1-state 1-condition to 2-state 1-condition) improved *R*_free_ (average improvement: 0.016 *±* 0.005). The improvement does not appear to depend on the temperature. For a 2-state model, increasing the number of conditions from 1 to 3 improved *R*_free_ in 5 of the 6 cases (average improvement: 0.035*±*0.018).

**Table 3:** Benchmark results. The *R*_free_ of the best-scoring 2-state model for each dataset when 1, 2, and 3 datasets are used in modeling. The best-scoring model for a dataset is the sample model that best satisfies the free reflections of a dataset. The same set of free reflections has been used for all datasets. The best-scoring model for a dataset could be informed by any combination of 1, 2, or 3 datasets that includes the dataset. The *R*_free_ of the (1-state) PDB model is also reported. The *R*_free_ of each PDB model is computed against a distinct set of withheld reflections.

| Method | Model Representation | | $R_{\text{free}}$ | | | | | |
| --- | --- | --- | --- | --- | --- | --- | --- | --- |
|  | States | Cond. | 100 K<br>1.55 Å | 240 K<br>1.53 Å | 277 K<br>2.19 Å | 298 K<br>1.88 Å | 298 K*<br>2.00 Å | 310 K<br>1.96 Å |
| MSMC | 1 | 1 | 0.323 | 0.316 | 0.350 | 0.335 | 0.351 | 0.349 |
| MSMC | 2 | 1 | 0.304 | 0.295 | 0.342 | 0.314 | 0.335 | 0.337 |
| MSMC | 2 | 2 | 0.305 | 0.299 | 0.328 | 0.304 | 0.315 | 0.315 |
| MSMC | 2 | 3 | 0.278 | 0.289 | 0.296 | 0.282 | 0.287 | 0.283 |

**Table 4:** Benchmark results. The corresponding *R*_work_ for all entries in previous table.

| Method | Model Representation | | $R_{\text{work}}$ | | | | | |
| --- | --- | --- | --- | --- | --- | --- | --- | --- |
|  | States | Cond. | 100 K<br>1.55 Å | 240 K<br>1.53 Å | 277 K<br>2.19 Å | 298 K<br>1.88 Å | 298 K*<br>2.00 Å | 310 K<br>1.96 Å |
| PDB | 1 | 1 | 0.182 | 0.160 | 0.199 | 0.191 | 0.195 | 0.198 |
| MSMC | 1 | 1 | 0.198 | 0.194 | 0.187 | 0.190 | 0.197 | 0.204 |
| MSMC | 2 | 1 | 0.187 | 0.180 | 0.176 | 0.176 | 0.180 | 0.192 |
| MSMC | 2 | 2 | 0.185 | 0.182 | 0.176 | 0.176 | 0.183 | 0.187 |
| MSMC | 2 | 3 | 0.190 | 0.188 | 0.173 | 0.184 | 0.177 | 0.191 |
|  | States | Cond. | 100 K<br>1.55 Å | 240 K<br>1.53 Å | 277 K<br>2.19 Å | 298 K<br>1.88 Å | 298 K*<br>2.00 Å | 310 K<br>1.96 Å |
| MSMC | 1 | 1 | 0.276 | 0.271 | 0.216 | 0.230 | 0.227 | 0.230 |
| MSMC | 2 | 1 | 0.253 | 0.245 | 0.199 | 0.198 | 0.210 | 0.215 |
| MSMC | 2 | 2 | 0.265 | 0.239 | 0.209 | 0.227 | 0.231 | 0.232 |
| MSMC | 2 | 3 | 0.240 | 0.237 | 0.207 | 0.209 | 0.208 | 0.209 |

**Figure 6.**
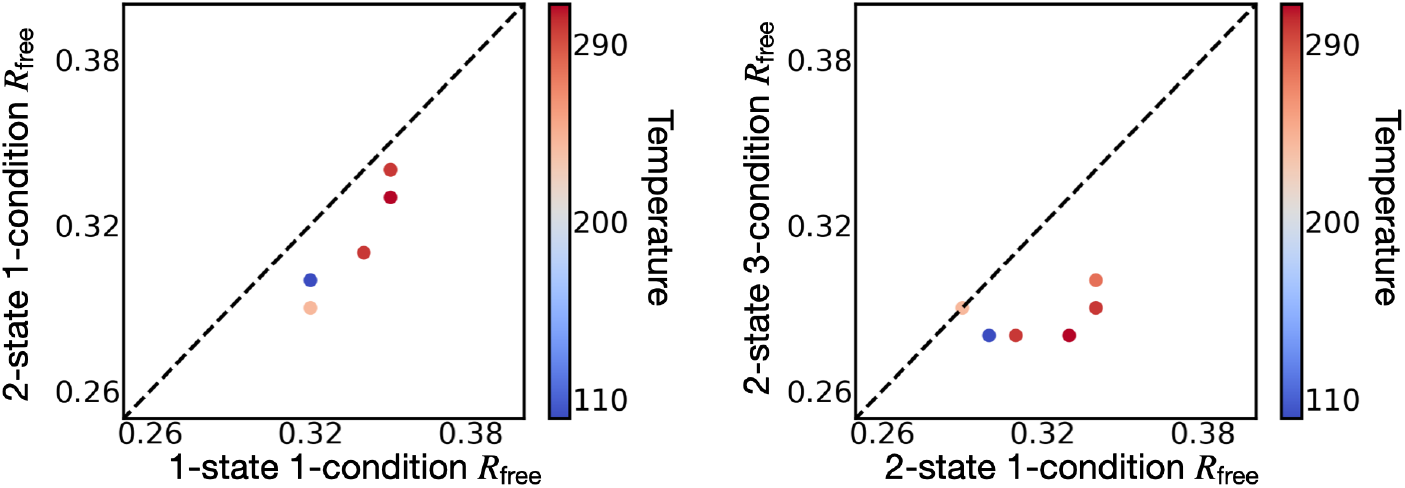
2-state multi-condition models of SARS-CoV-2 Mpro. Left, the *R*_free_ of the best-scoring 2-state multi-condition model *vs* the *R*_free_ of the PDB-deposited model for each dataset (6 in total), colored by temperature. Right, the *R*_free_ of the best-scoring 2-state 2-condition model *vs* the *R*_free_ of the best-scoring 2-state 1-condition model, colored by temperature. In both cases, a point below the line indicates improvement for the corresponding dataset.

In contrast to adding an additional state, the *R*_free_ improvement from additional conditions for a dataset appears to depend on temperature. The average improvement for 100K and 240K was 0.016 while the improvement of 277K, 298K, 298K*, and 310K was 0.045. This improvement may result from smaller differences between the crystals at 277K, 298K, 298K*, and 310K, as well as a higher importance of additional information for fitting models into lower quality datasets.

The PDB models [Jin+20] were refined with *Phenix*.*refine*, enhancing fit to the experimental data by refining atomic positions, isotropic atomic B-factors, atomic occupancies, placement of ligands/solvent, *etc*. We have not yet implemented a comparable protocol for refining multi-state multi-condition models; as mentioned above, our current software produces models without solvent and ligand molecules, with constant isotropic atomic B-factors, and with constant atomic occupancies. To estimate the scope for improving our multi-state multi-condition models further by including additional variables into model representation and more exhaustive sampling of existing variables, we used *Phenix*.*refine* protocol to refine individual states in our multi-state multi-condition models. Thus, this protocol refined all atomic positions and B-factors of all states, placed solvent atoms, and included a Zn+ ligand found in all PDB-deposited structures, albeit effectively for a single state at a time (Tables 5, 8). In all cases, *Phenix*.*refine* improves *R*_free_ of the starting model (average improvement: 0.051 *±* 0.010). Therefore, we suggest that multi-state multi-condition modeling can be improved further by adding additional variables to model representation and perhaps more exhaustive sampling, following *Phenix*.*refine*.

**Table 5:** Benchmark results. The *R*_free_ of the best-scoring 2-state model for each dataset when 1, 2, and 3 datasets are used in modeling. The best-scoring model for a dataset is the sample model that best satisfies the free reflections of a dataset. The same set of free reflections has been used for all datasets. The best-scoring model for a dataset could be informed by any combination of 1, 2, or 3 datasets that includes the dataset. The *R*_free_ of the (1-state) PDB model is also reported. The *R*_free_ of each PDB model is computed against a distinct set of withheld reflections.

| Method | Model Representation | | $R_{\text{free}}$ | | | | | |
| --- | --- | --- | --- | --- | --- | --- | --- | --- |
|  | States | Cond. | 100 K<br>1.55 Å | 240 K<br>1.53 Å | 277 K<br>2.19 Å | 298 K<br>1.88 Å | 298 K*<br>2.00 Å | 310 K<br>1.96 Å |
| PDB | 1 | 1 | 0.224 | 0.205 | 0.253 | 0.228 | 0.240 | 0.247 |
| MSMC | 1 | 1 | 0.228 | 0.231 | 0.241 | 0.234 | 0.249 | 0.254 |
| MSMC | 2 | 1 | 0.223 | 0.226 | 0.251 | 0.235 | 0.249 | 0.258 |
| MSMC | 2 | 2 | 0.221 | 0.223 | 0.243 | 0.227 | 0.239 | 0.247 |
| MSMC | 2 | 3 | 0.227 | 0.228 | 0.237 | 0.230 | 0.239 | 0.249 |

## 5 Discussion

We developed and evaluated a Bayesian method to compute a model depicting multiple conformational states (multi-state model) from multiple X-ray datasets representing distinct experimental conditions (multi-condition datasets). Next, we discuss, in turn, advantages and challenges of computing such models and their potential applications.

### 5.1 Using a multi-state model representation

We highlight a key advantage and four current disadvantages of our multi-state representation.

The key advantage of using a multi-state representation is that it is in principle a more faithful depiction of a crystal than a single-state representation: X-ray scattering is a function of positions and types of all atoms in the crystal; thus, an accurate scoring function requires a model representation that includes all atomic positions in the unit cell. The global minimum of such a scoring function can then be closer to the native than the global minimum of a scoring function that is less reflective of the actual content of the crystal. In general, the set of possible atomic positions that need to be considered is large, because of the compositional and structural heterogeneity, including multiple states of a protein, solvent molecules, and ligands. Here, we addressed the conformational heterogeneity of the protein by explicitly depicting it with a set of several discrete conformational states, each one corresponding to a mode of a smooth distribution of conformations in the crystal. When the optimal representation is unknown before modeling, the number of states may be enumerated from 1 to the maximal computationally feasible number of states, followed by applying a model selection criterion, such as the Bayesian or Akaike Information Criterion, to select the final model [SS04]; such a criterion would select the smallest number of states that is sufficient to explain the X-ray data within its uncertainty.

The first disadvantage of using a multi-state representation is the difficulty of interpreting the multi-state model because X-ray data are degenerate with regard to assigning atomic positions to specific states in a multi-state representation. A multi-state scoring function evaluates the collective satisfaction of an X-ray dataset by all states. In other words, it assess how well all states together satisfy the data, not how well any single state in the set of states satisfies the data. In X-ray crystallography, a 1-state scoring function can be generalized to a multi-state scoring function by exploiting the equivalence between the forward model for a weighted set of states and the weighted sum of single state forward models (Figure 2). However, the corresponding degeneracy of the X-ray data results in uncertainty of assigning an atom position to a particular state, in turn complicating interpretation of a multi-state model: while a multi-state model that perfectly satisfies the X-ray data specifies the atomic positions in all states correctly, it should not be interpreted as assigning these positions to any particular state. To resolve this assignment ambiguity, additional information, such as a molecular mechanics force field, must be included in the scoring function.

The second disadvantage is that sampling of single-state models is more challenging, due to the higher dimensionality and potential ruggedness of the scoring function landscape. It is conceivable that the smoothness of the landscape would be increased by re-formulating it without some nuisance parameters, such as the scaling factors *α* and *β*. The model search may be further improved using more advanced sampling algorithms, such as simulated annealing [KGV83], instead of the constant-temperature molecular dynamics simulations used here. Additionally, future sampling may benefit from including additional information about protein stereochemistry (*eg*, sampling only known rotameric side chain states). The third disadvantage is that the multi-state representation may introduce more variables than necessary to accurately depict the conformational heterogeneity found in a protein crystal. For example, the positions of atoms in solvent exposed side chains and loops tend to vary more than those in the core ([WB00]). However, a representation containing multiple states will depict both static and variable regions with an equal number of variables. Increasing the number of variables increases the computational cost of sampling and thus the likelihood that the models overfits the data used for modeling. This challenge may be addressed by employing a hierarchical representation that only introduces structural variables, such as multiple backbone conformations and sidechain rotamers, where they are needed ([WF24; Ril+21]). Another solution may be including restraints that maximize the similarity among the model states, as long as the data are satisfied sufficiently.

The fourth disadvantage is that our method does not include solvent and other non-protein atoms in the sampling of multi-state models. Even though we use *Phenix*.*refine* to build and refine a single copy of non-protein atoms into the model, we suspect that a method that can build in and refine partially occupied non-protein atoms for each specific state may significantly improve the *R*_free_.

### 5.2 Using multiple X-ray datasets

In our method, multiple X-ray datasets are incorporated into modeling *via* the scoring function. We highlight a key advantage and two current disadvantages of using multiple X-ray datasets.

The key advantage of a multi-condition scoring function is that it considers more information than a single-condition scoring function. Benchmarking shows that models informed by 2 X-ray datasets satisfy the data better than models informed by 1 X-ray dataset, as measured by R-free, for all synthetic and experimental datasets, regardless of the number of states (Table 1, Table 2). Two X-ray datasets may yield a more accurate model than a single dataset because the scoring function is more accurate and/or the scoring function landscape can be sampled better, as follows.

First, we consider why a multi-condition scoring function may be more accurate, in the sense that its global minimum is closer to the native. The increased accuracy may result from lower random and systematic errors. The random error of the scoring function generally results from the random error in the data. In contrast, the systematic error of the scoring function can be seen as resulting from its incorrect formulation, an error in model representation, and/or systematic error in the data. We initially hypothesized that using multiple X-ray datasets will decrease the random error of the scoring function (*cf*, the standard error of the mean is equal to the standard deviation divided by the square root of the number of samples). However, scoring models against synthetic reflections with a large random error (*σ* = 5%) and no systematic error results in a highly accurate scoring function, regardless of the number of conditions (*ie*, there is no decoy model that better satisfies the data than the native model) (Figure 3), due to the scoring function for a single X-ray dataset already depending on thousands of independent reflections. Thus, we conclude that the error found in models computed from one or more X-ray datasets is primarily a reflection of the systematic errors in the scoring function and imperfect sampling of the scoring function. The systematic error likely originates from an imperfect depiction of the crystal and unaccounted experimental errors. For a given model representation, these sources of systematic error likely do not depend significantly on the number of datasets. We therefore conclude that the improvement in modeling from including multiple conditions must be attributed to improved sampling of the scoring function, contrary to our initial hypothesis.

Second, we consider why a multi-condition scoring function may be sampled more easily than a single-condition scoring function. In addition to the global minimum, the scoring function has a large number of local minima (models corresponding to these local minima may increase the satisfaction of the reflections included in modeling without a commensurate increase in the satisfaction of the reflections not included in modeling). We expect that the states corresponding to the global minimum of the scoring function based on each individual dataset are similar, while their local minima are less conserved. Thus, a multi-condition scoring function is likely smoother than any single-condition scoring function, thus making sampling more efficient. This hypothesis is consistent with the observed improvements in R-free upon adding a second dataset: These improvements are larger for lower resolution datasets with lower data-to-parameter ratios (277 K, 298 K, 298 K*, and 310 K) than for the higher resolution datasets (100 K and 240 K) (Table 2). Thus, our method may be especially useful for computing multi-state models from lower resolution X-ray datasets, commonly produced from higher temperature experiments.

The first disadvantage of a multi-condition scoring function results from a potential increase in the number of states as the number of experimental conditions increases, in turn increasing the risk of overfitting and insufficient sampling. In our benchmark, for example, using 3 instead of 2 datasets tends to decrease the accuracy of the 2-state and 3-state models (Table 2).

The second disadvantage is the increase in the number of model variables, corresponding to an additional set of state weights for each condition, again increasing the risk of overfitting and insufficient sampling. Accurately fitting the matrix of weights of a multi-state multi-condition model remains a significant challenge. Currently, our approach samples weights randomly, relying on parallel molecular dynamics simulations. Further development is required to more efficiently sample the weight variables.

### 5.3 Applications of multi-condition multi-state modeling

Bayesian multi-state multi-condition modeling may have several applications in X-ray crystallography. First, in serial crystallography, a series of small and often imperfect crystals are exposed to the X-ray beam, generating partial diffraction patterns at a series of time points [Bar+22]. This trajectory could be represented as differently weighted mixtures of the same states along the sampled time points. Our multi-state multi-condition scoring function would inform a model of each state at each time point based on the totality of the X-ray data, potentially improving the accuracy of the models as demonstrated here. Second, in a screen of many fragments, the protein is crystallized in the presence of one or a few fragments at a time, corresponding to a single condition [KFD20]. Fragments often have low affinity, and therefore only a fraction of the protein molecules in the crystal bind the fragments. The multi-state multi-condition representation can conceivably combine data from multiple experiments in several ways. For example, a multi-state multi-condition model may be computed where the states correspond to each fragment”s apo state and the dominant holo state. As a result, X-ray diffraction patterns from each fragment will inform all states. Third, non-Bragg scattering is often ignored in modeling based on X-ray crystallography data, resulting in the loss of information that could potentially be useful in modeling [Van+16]. For example, diffuse scattering contains information on the correlated motions of atoms in the crystal [Pei+23]. Our Bayesian framework is in principle poised to include another likelihood term based on diffuse scattering data, to compute a more accurate and precise multi-state model.

Finally, to facilitate the application of our method by scientists to many systems, we implemented it as a module of our open-source *Integrative Modeling Platform* (IMP) software [Rus+12], freely available at https://integrativemodeling.org/2.20.2/doc/manual/. For the calculations described here, IMP relies critically on Phenix [**ref**], highlighting the need for open-source and modular software packages [Han+22].

## 6 Acknowledgments

We are grateful to Dr. Benjamin M. Webb for help with the IMP software, and Nebius Inc. for their support of IMP software development (BWB is Nebius Fellow). We also acknowledge NIH grants R01GM083960 and U19AI171110 (AS) as well as R35GM145238 (JSF).

## Notes

### Competing Interest Statement

The authors have declared no competing interest.

